# GHIST 2024: The 1st Genomic History Inference Strategies Tournament

**DOI:** 10.1101/2025.08.05.668560

**Authors:** Travis J. Struck, Andrew H. Vaughn, Austin Daigle, Dylan D. Ray, Ekaterina Noskova, Jaison J. Sequeira, Svetlana Antonets, Elizaveta Alekseevskaya, Elizaveta Grigoreva, Evgenii Raines, Eilish S. McMaster, Toby G. L. Kovacs, Aaron P. Ragsdale, Andrés Moreno-Estrada, Katie E. Lotterhos, Adam Siepel, Ryan N. Gutenkunst

**Author notes:** These authors contributed equally to this work (Travis J. Struck, Andrew H. Vaughn).

## Abstract

Evaluating population genetic inference methods is challenging due to the complexity of evolutionary histories, potential model misspecification, and unconscious biases in self-assessment. The Genomic History Inference Strategies Tournament (GHIST) is a community-driven competition designed to evaluate methods for inferring evolutionary history from population genomic data. The inaugural GHIST competition ran from July to November 2024 and featured four demographic history inference challenges of varying complexity: a bottleneck model, a split with isolation model, a secondary contact model with demographic complexity, and an archaic admixture model. Data were provided as error-free VCF files, and participants submitted numerical parameter estimates that were scored by relative root mean squared error. Approximately 60 participants competed, using diverse approaches. Results revealed the current dominance of methods based on site frequency spectra, while highlighting the advantages of flexible model-building approaches for complex demographic histories. We discuss insights regarding the competition and outline the next iteration, which is ongoing with expanded challenge diversity. By providing standardized benchmarks and highlighting areas for improvement, GHIST represents a substantial step toward more reliable inference of evolutionary history from genomic data.

---

Population genetic inference aims to reconstruct the recent evolutionary history of populations from genomic variation data. This field has seen explosive growth, driven by the increasing availability of whole-genome sequencing data from diverse groups of humans and other species (Pool et al. 2010). But population genetic inference is inherently challenging. First, the stochasticity of the evolutionary process means that the same history can produce different genetic patterns. Second, different histories can produce similar patterns of genetic variation, creating an identifiability problem (Myers et al. 2008; Lapierre et al. 2017; Lawson et al. 2018; Rosen et al. 2018). Third, real populations rarely conform to the simplified models typically used for inference, leading to potential biases when models are misspecified (Loog 2021; Momigliano et al. 2021). Finally, computational constraints often necessitate approximations that may impact accuracy.

Many methods for population genetic inference exist. For example, site frequency spectrum (SFS) methods examine the distribution of allele frequencies within and among populations (Marth et al. 2004; Gutenkunst et al. 2009; Excoffier et al. 2013). Linkage-based approaches analyze patterns of linkage disequilibrium or identity-by-descent (IBD) segments (Harris & Nielsen 2013; Browning & Browning 2015). Markovian coalescent methods reconstruct recent genealogical relationships among samples (Li & Durbin 2011; Schiffels & Durbin 2014), while ancestral recombination graph (ARG) methods explicitly reconstruct the genealogical history including recombination events (Rasmussen et al. 2014; Kelleher et al. 2019; Speidel et al. 2019). More recently, machine learning approaches apply supervised learning to haplotype matrices or summary statistics (Schrider & Kern 2018; Flagel et al. 2019; Sanchez et al. 2021; Tran et al. 2024). Each approach captures only a portion of the information contained in genomic data, and different methods excel in different scenarios.

Papers describing new inference methods typically benchmark against existing approaches, but these self-assessments are often biased (Norel et al. 2011; Boulesteix 2015), if unconciously. First, method developers naturally focus on scenarios where their approaches excel, potentially masking weaknesses. Second, developers have intimate knowledge of optimal parameter settings for their own methods but may use default parameters for competing methods, leading to unfair comparisons. Lastly, developers benchmarking their own tools know the ground truth they simulated, enabling unconscious bias toward that truth. Best-practice guidelines for benchmarking studies (Boulesteix 2015; Lotterhos et al. 2022) can reduce, but not eliminate, these biases.

Independent benchmarking studies can provide more reliable conclusions than developer-driven benchmarking (Boulesteix et al. 2013), and they have been conducted in population genomics, but limitations remain. While developing a data simulation framework for the community, the stdpopsim project compared methods for inferring demographic history, distributions of fitness effects, and selective sweeps, although not systematically (Adrion et al. 2020; Gower et al. 2025). For demographic history inference, parametric SFS-based methods have been compared with non-parametric SFS-based (Lapierre et al. 2017) and Markovian coalescent methods (Beichman et al. 2017). The confounding effects of background selection on such inference have been studied for SFS-based and Markovian coalescent methods (Johri et al. 2021) and ARG-based methods (Marsh & Johri 2024). Brandt et al. (2022) evaluated the accuracy of ARG inference methods in estimating coalescence times, Peng et al. (2025) evaluated ARG-based methods for predicting historical polygenic scores, and Patton et al. (2019) evaluated non-parametric methods for demographic history inference under varying genome assembly quality. Although these studies have investigated many different tools, each has been carried out by a small group of authors, and their expertise in the tools tested can strongly influence benchmark results (Lotterhos et al. 2016; Weber et al. 2019). And because each of these studies is singular, it is difficult to assess progress in the field from them.

Community-based competitions have proven effective at driving innovation across multiple domains of computational biology (Meyer et al. 2011). The Critical Assessment of Protein Structure Prediction (CASP), running since 1994, is perhaps the most successful (Moult et al. 1995). By providing semi-annual blind tests of protein structure prediction methods, CASP has catalyzed remarkable improvements, culminating in the 14th competition with AlphaFold 2’s breakthrough performance that revolutionized structural biology (Jumper et al. 2021; Kryshtafovych et al. 2021). Similarly, challenges from DREAM (Dialogue for Reverse Engineering Assessment and Methods) have addressed diverse problems in systems biology and genomics, from gene regulatory network inference to disease prediction (Stolovitzky et al. 2007;

Marbach et al. 2012; Saez-Rodriguez et al. 2016). More recently, the Critical Assessment of Genome Interpretation focuses on predicting phenotypic consequences of genetic variants, driving improvements in variant effect prediction (Critical Assessment of Genome Interpretation Consortium 2024). PrecisionFDA challenges evaluate methods for variant calling, genome assembly, and other genomics tasks, setting standards for precision medicine applications (Olson et al. 2022). These examples illustrate the power of competition-based assessments of computational biology methods. In evolutionary inference, real-world data for which the truth is known is typically lacking (see Randall et al. (2016) for an exception), but modern simulators capture enough features of real data to provide valuable insights (Baumdicker et al. 2022; Haller & Messer 2023).

The Genomic History Inference Strategies Tournament (GHIST, pronounced /gIst) adapts the successful competition model to address the specific challenges of population genetic inference. Here, we report results from the first competition, which consisted of four challenges focused on inferring demographic history. The competition attracted many participants, demonstrated the feasibility of the model, and revealed current community practices.

## Methods

The organizing team (authors Struck and Gutenkunst) and the design committee (authors Lotterhos, Moreno-Estrada, Ralph, and Siepel) collaborated closely to develop the structure of the first GHIST competition. Although creating highly complex challenges was tempting, we prioritized accessibility to ensure early community engagement and success. We chose demographic history inference as the competition’s focus, because it is foundational to many other population genetic analyses, it allows comparison across a variety of established methods, and aligns with the organizers’ expertise. To further encourage participation, we outlined a proactive communication strategy and offered authorship on the resulting paper as recognition for the top-performing competitors.

The structure of the first GHIST competition was developed in collaboration between the organizing team (authors Struck and Gutenkunst) and design committee (authors Lotterhos, Moreno-Estrada, Ralph, and Siepel). While it was tempting to develop extremely complex challenges, accessibility was deemed important for early success of the competition. It was decided to focus on demographic history inference, because it is foundational for other population genetics inference tasks, there are many methods to compare, and because the organizers have specific expertise. To engage community participation, a preliminary plan for communications was also developed. Lastly, to incentivize participation, it was decided that top competitors would be offered authorship on the resulting paper.

The inaugural GHIST competition consisted of four demographic history inference challenges. These were a simple bottleneck, a simple split with migration, a complex split with secondary contact, and a complex archaic admixture scenario. Competitors could submit to any challenge(s) they chose, in any order. The scenarios were parameterized such that existing methods were expected to have good statistical power and sample sizes were set to be similar to contemporary non-human data sets. For all four challenges, the data were simulated using the WrightFisher coalescent method msprime (Baumdicker et al. 2022) and distributed as error-free Variant Call Format (VCF) files (Danecek et al. 2011), with only biallelic sites and correct ancestral states provided. To minimize complexity, mutation and recombination rates were uniform across the simulated regions, and selection was absent.

For each challenge, competitors reported estimates for a small number of key population genetic parameters, such as population sizes, divergence times, or admixture proportions. They were told the total size of the simulated region and the true simulated mutation and recombination rates. Entries were scored based on the relative root-mean-squared error between submitted 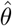 and true parameter values *θ*:

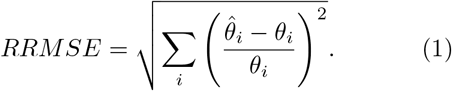

This interpretable metric allowed comparison across parameters of different scales and penalized both over- and under-estimation equally. For each challenge, the leaderboard was ranked based on RRMSE scores, with lower scores indicating better performance. To allow methodological refinement, competitors were allowed five submissions for each challenge. In addition to their inferences, competitors were asked to submit a brief write-up of their approach, including software tools used and the logical flow of their analyses. The scripts for generating the data and scoring submissions are available at https://github.com/tjstruck/GHIST-2024-paper.

The competition was hosted on the Synapse platform developed by Sage Bionetworks, a not-for-profit organization that promotes open science and collaborative research. Synapse provided automated handling of competitor submissions, including timestamps, versioning, validation, and real-time leaderboards. The integrated wiki functionality was used for competition documentation and tutorials, and discussion boards enabled competitors to ask questions of the organizers. The Synapse site for the first GHIST competition is available at https://synapse.org/Synapse:syn51614781, and the main GHIST website is at https://ghi.st.

The inaugural GHIST competition ran from July to November 2024, to span the summer conference season and the beginning of the academic term (Fig. 1). It began with a kickoff workshop at the Society for Molecular Biology and Evolution (SMBE) conference in Puerto Vallarta, Mexico, where participants were introduced to the competition, analyzed data from the Bottleneck challenge using dadi-cli (Huang et al. 2023), and submitted their inferences. The competition extended into the academic term to enable new students to participate as a training opportunity. The competition was promoted in-person at the SMBE and Evolution conferences, through posts to the Evoldir, dadi user, and fastsimcoal user mailing lists, and through targeted emails to specific investigators known to the organizers. It was also promoted through posts on X and Bluesky by the organizers and SMBE.

**Figure 1.**
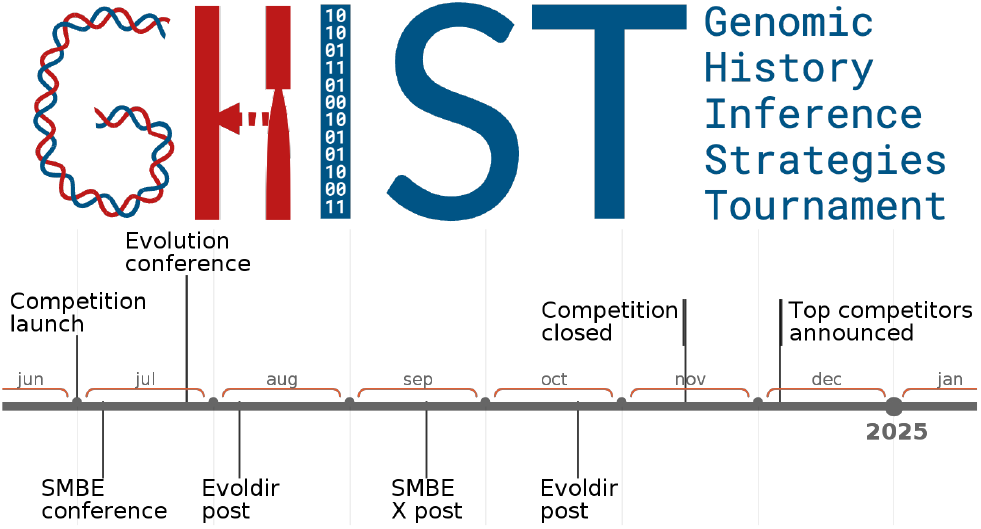
Timeline of the first GHIST competition, including notable promotion events.

## Results

The inaugural GHIST competition attracted approximately 60 participants spanning career stages from graduate students to senior faculty. Participation varied across challenges, with more entries for the simpler challenges. Competitors employed a variety of approaches, with top competitors mostly relying on the site frequency spectrum (SFS). A variety of software was employed, including custom pipelines.

### Bottleneck challenge

The first challenge involved a simple bottleneck (Fig. 2A), with competitors inferring the timing and magnitude of the population decline. Competitors were given 100 megabases (Mb) of data from 20 diploid individuals, yielding 219 thousand biallelic variants.

**Figure 2.**
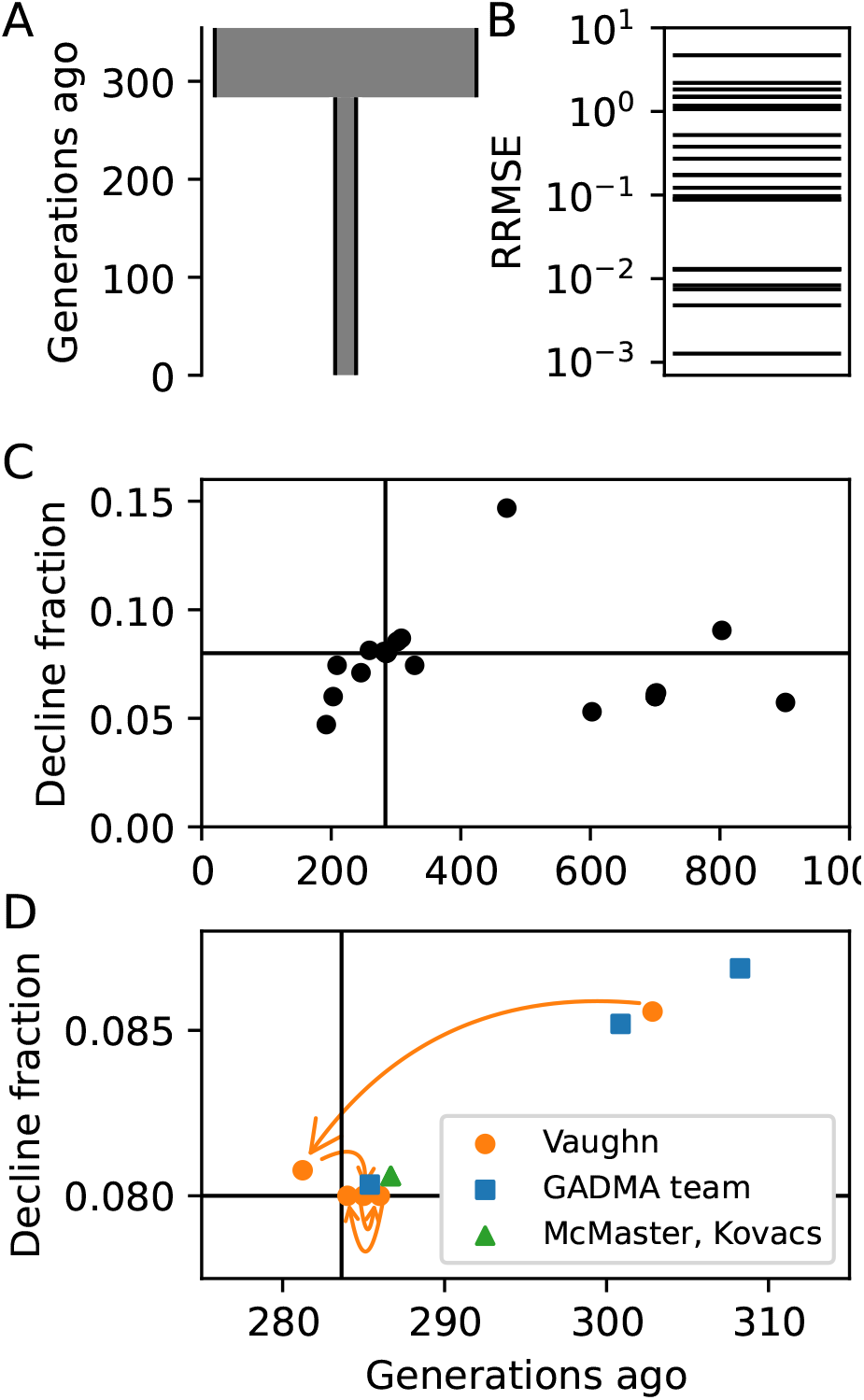
GHIST 2024 Bottleneck challenge. A) True simulated demographic history. B) Relative root mean square error scores of submissions. C) Parameter inferences of majority of submissions. True values are indicated by solid lines. D) Parameter inferences zoomed close to true values to indicate top competitors. Arrows indicate competitor Vaughn’s leaderboard optimization procedure.

Submissions for the bottleneck challenge showed a range of strategies and accuracy. The RRMSE values of submissions spanned orders of magnitude (Fig. 2B). Almost all submissions successfully identified the presence of a bottleneck (Fig. 2C), but only a few were highly accurate.

The most accurate submissions for the Bottleneck challenge used site frequency spectrum (SFS) approaches (Fig. 2D). Competitor Vaughn developed a custom approach using an analytic result for piecewise constant demographic histories to calculate the expected model SFS (DeWitt et al. 2021) and the Kullback-Leibler (KL) divergence to measure differences between model and data spectra. Competitors McMaster and Kovacs used the SFS-based methods dadi-cli (Huang et al. 2023) and fastsimcoal2 (Excoffier et al. 2021) and the Markovian coalescent tool SMC++ (Terhorst et al. 2017) for their submissions. Competitor Noskova led a team using her GADMA (Noskova et al. 2020, 2023) framework, using the dadi (Gutenkunst et al. 2009), moments (Jouganous et al. 2017), and momi2 (Kamm et al. 2020) engines for calculating model spectra.

A surprise was that top competitor Vaughn metagamed the challenge by using the leaderboard to optimize his submissions. He made an excellent first submission (Fig. 2D) based on the provided data, but this would not have been enough to win the challenge. To improve his result, he correctly deduced that the challenge simulation used round parameter values, and he used his remaining four submissions to search through the parameter space using the leader-board RRMSE score to converge on nearly the exact values.

### Split with Isolation challenge

The second challenge involved two populations that diverged without subsequent gene flow, representing geographic isolation (Fig. 3), with competitors inferring the contemporary population sizes and the timing of the split. They were given 100 Mb of data from 22 and 18 individuals from the two populations, yielding 1.2 million biallelic variants.

**Figure 3.**
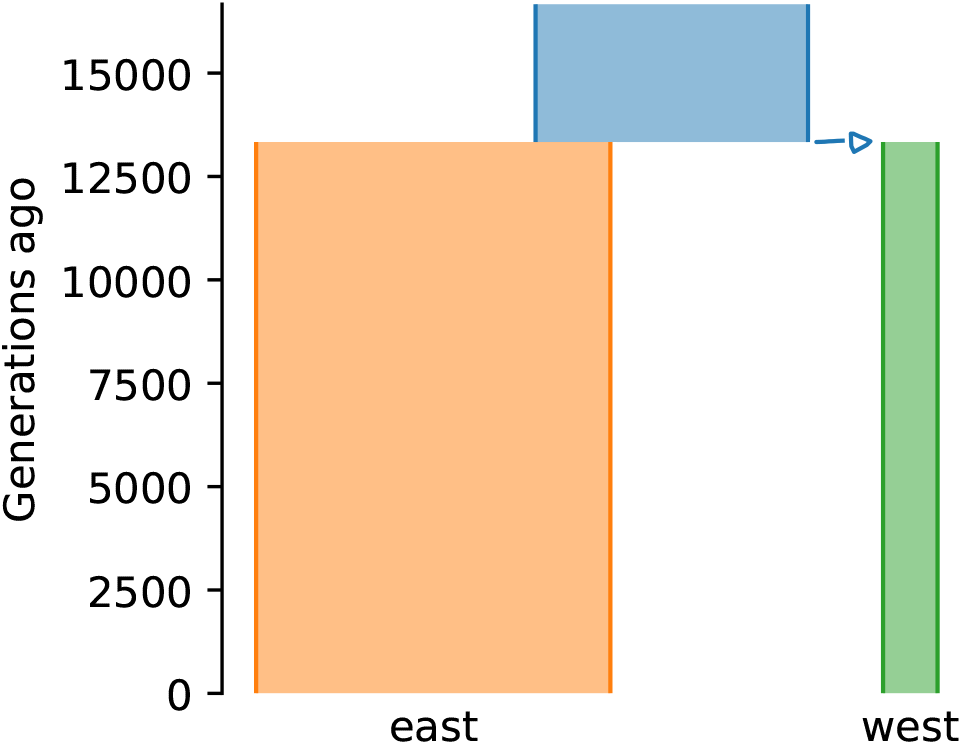
True demographic history for the Split with Isolation challenge, represented using demesdraw (Gower et al. 2022).

Performance on this challenge was generally strong, with several competitors achieving high accuracy for all three parameters (Table 1). SFS-based methods performed well in this challenge, with competitor Vaughn using tskit (Kelleher et al. 2016; Wong et al. 2024) to calculate expected spectra and KL divergence to fit the model and McMaster and Kovacs using dadi-cli for inference. The Team of Daigle and Ray used a machine learning approach. They first used dadi to identify the relevant ranges of parameter values, then simulated data over those ranges with msprime, and then used scikit-allel (Miles et al. 2024) and pylibseq (Thornton 2003) to calculate summary statistics, including statistics based on the SFS, haplotypes, and LD decay. These summary statistics were then passed to a multi-layer perceptron for inference. However, their best-scoring submission for this challenge simply employed dadi. As in the first Challenge, competitor Vaughn achieved the top score by strategically rounding his inferences.

**Table 1:** Top submissions for the Split with Isolation challenge.

| $RRMSE$ | $N_{east}$ | $N_{west}$ | $T$ | Competitor | Approach |
| --- | --- | --- | --- | --- | --- |
| truth | 130000 | 20000 | 13333 |  |  |
| 0.005 | 130000 | 20100 | 13300 | Vaughn | metagaming |
| 0.027 | 126686 | 19973 | 13233 | McMaster, Kovacs | dadi-cli |
| 0.029 | 126941 | 19867 | 13119 | Daigle, Ray | dadi |
| 0.031 | 125992 | 20109 | 13301 | Vaughn | tskit SFS |

### Secondary Contact challenge

The third challenge involved secondary contact between isolated populations, with complexity in population size histories that no parametric model was expected to capture (Fig. 4A). Competitors were tasked with inferring the contemporary population sizes, timing of the split and recontact, and the rate of migration after recontact. Competitors were again given 100 Mb of data, from 22 diploid mainland individuals and 8 island individuals, for a total of 842 thousand biallelic sites.

**Figure 4.**
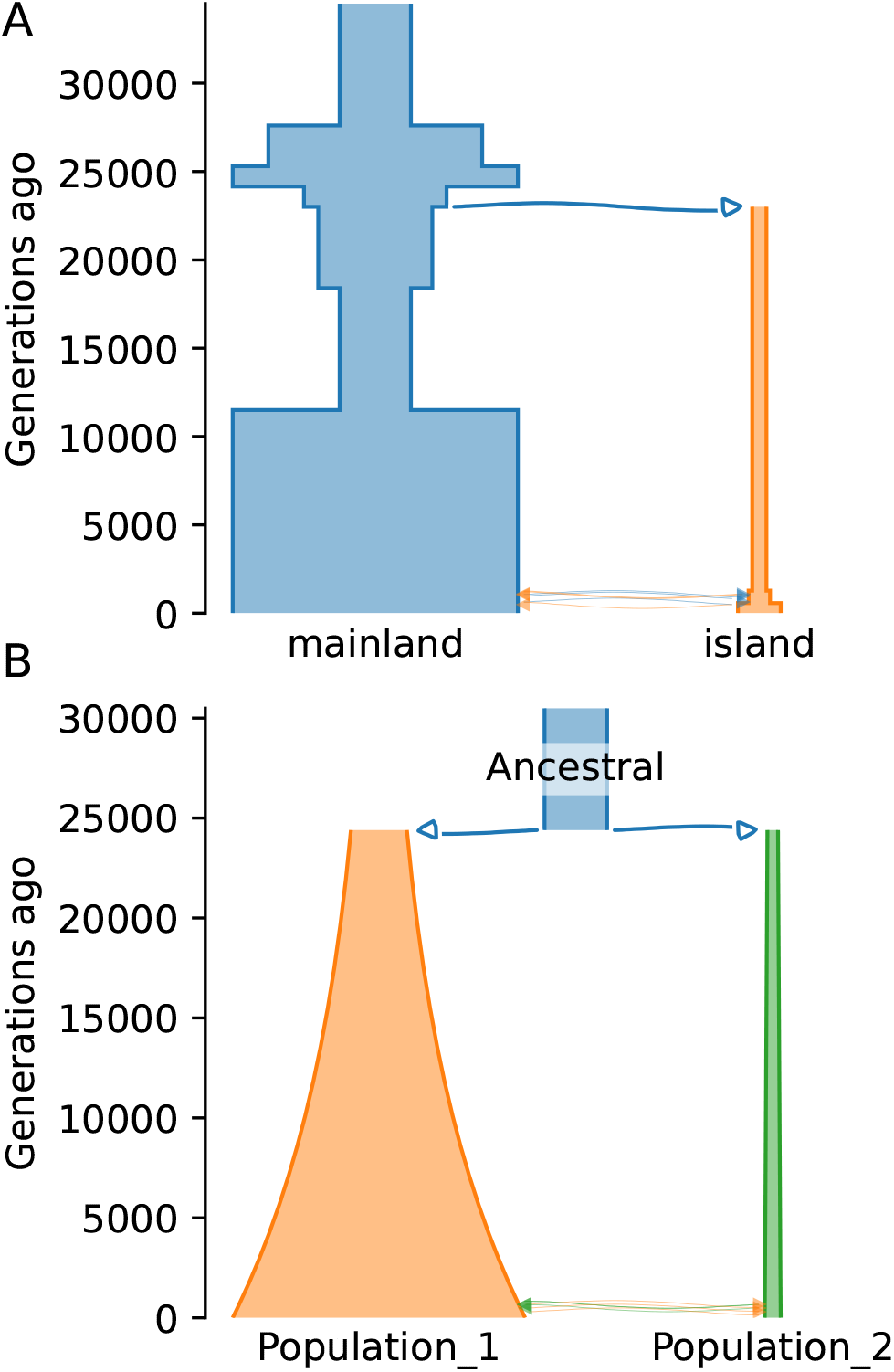
GHIST 2024 Secondary Contact challenge. A: True demographic history, represented using demesdraw (Gower et al. 2022). B: Top-scoring model, from the GADMA team.

As expected, this challenge was more difficult than the previous two, with no submission accurately estimating all parameters (Table 2). The team of Daigle and Ray did well with their machine learning approach based on summary statistics. Competitor Vaughn’s top submissions were all based on leaderboard optimization after his initial inference. All these submissions assumed simple constant population size histories, like the truth in the Split with Isolation challenge. The best performance came from the GADMA team, using the moments engine (Table 2). GADMA automatically builds and refines models of increasing complexity, and their best model allowed for growth in both populations (Fig. 4B), perhaps enabling their model to account for some of the effects of the true complex population size changes.

**Table 2:** Top submissions for Secondary Contact challenge.

| $RRMSE$ | $N_{\text{main}}$ | $N_{\text{island}}$ | $T_{\text{split}}$ | $T_{\text{mig}}$ | $m$ | Competitor | Approach |
| --- | --- | --- | --- | --- | --- | --- | --- |
| truth | 240000 | 36000 | 23000 | 1277 | 5.0 |  |  |
| 0.92 | 284284 | 15567 | 24397 | 950 | 8.2 | GADMA team | GADMA w/ moments |
| 0.97 | 150000 | 35000 | 30000 | 200 | 5.0 | Vaughn | metagaming |
| 1.05 | 200000 | 12000 | 18000 | 310 | 5.0 | Vaughn | metagaming |
| 1.05 | 220000 | 60000 | 17000 | 300 | 5.0 | Vaughn | metagaming |
| 1.30 | 211999 | 10945 | 15818 | 215 | 1.8 | Daigle, Ray | summary stats perceptron |

### Archaic Admixture challenge

To probe a distinct but related form of inference, the final challenge involved archaic admixture. Competitors were tasked with inferring the timing and magnitude of admixture into two modern populations (Fig. 5). They were given 250 Mb of data from 20 and 16 samples for the modern populations, along with 1 to 3 samples from each of the potential archaic contributors, sampled 17,500 to 100,000 simulated years ago, for a total of 1.7 million biallelic sites.

**Figure 5.**
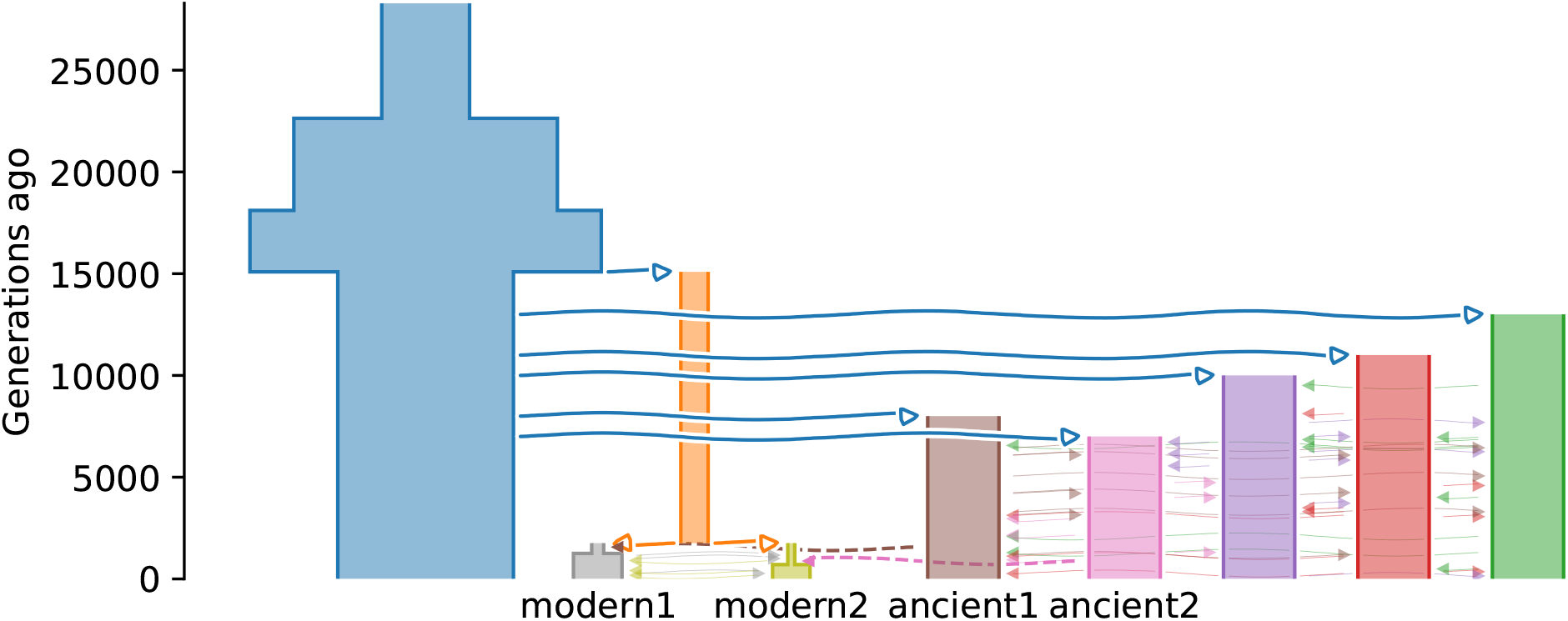
: True demographic history for Archaic Admixture challenge, represented using demesdraw (Gower et al. 2022).

The top competitors accurately estimated admixture proportions but were less accurate when estimating timings (Table 3). For each modern population, competitor Vaughn used tskit simulations to simulate a two-population model with archaic admixture from a ghost population and fit that to the SFS from the modern population. He then optimized the leader-board to refine his estimates. The GADMA software does not support ancient samples, so it could not be applied to this challenge. But the GADMA team used momi2 (Kamm et al. 2020) directly to fit models involving all five sample groups, achieving superior accuracy before metagaming.

**Table 3:**
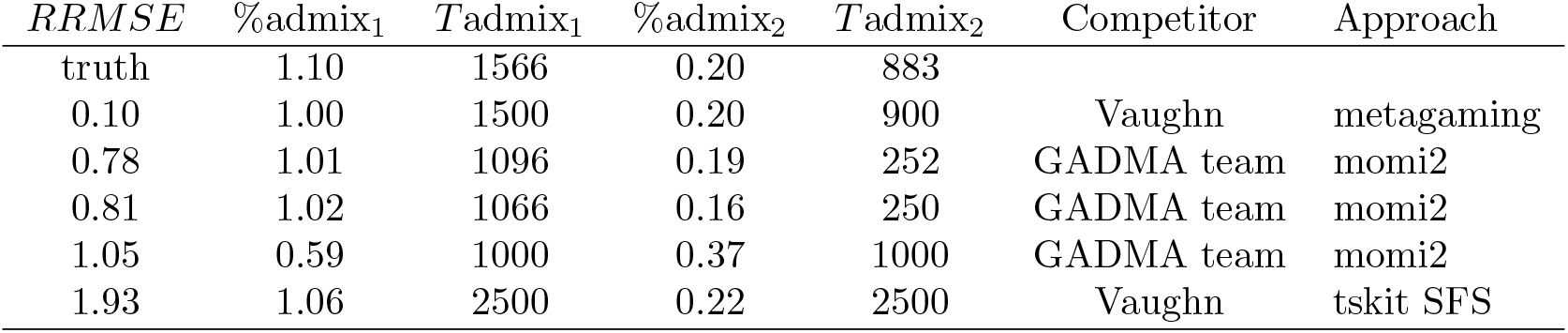
Top submissions for Archaic Admixture challenge.

## Discussion

The inaugural GHIST competition demonstrated the feasibility and value of a community-driven evaluation framework for population genetic inference methods. The Synapse platform proved robust and capable, and the range of challenges enabled accessibility while pushing the limits of existing inference methods. The conference-based launch and extended timeframe facilitated participation from diverse researchers, including students.

The GHIST competition provided several insights into the relative performance of inference approaches. Approaches based on the site frequency spectrum were most common and successful, because they are both accessible from established software tools and powerful for demographic inference. For the Bottleneck, Split with Isolation, and Secondary Contact challenges, the GADMA team directly compared SFS-based engines with the moments.LD engine that uses multi-population linkage disequilibrium statistics (Ragsdale & Gravel 2019, 2020), achieving better scores with SFS-based engines. Approaches based on machine learning showed promise but were not widely used by competitors. As those approaches become more accessible, we expect their representation and success to increase. Almost all approaches applied assumed prespecified parametric models, which may not capture the complexity of real demographic histories (Loog 2021). The exception was GADMA, and its success in the Secondary Contact challenge (Fig. 4A), which was designed to violate typical prespecification, highlights the importance of model flexibility when dealing with complex histories.

There were notable gaps in the methods employed by participants. Approaches based on ancestral recombination graphs show great promise for population genetics (Rasmussen et al. 2014; Kelleher et al. 2019; Speidel et al. 2019; Deng et al. 2024), but they were not applied to this competition, perhaps because of their high computational cost or complexity. The Archaic Admixture challenge (Fig. 5) was designed to encourage the use of specialized methods based on lengths of admixture tracts (Pool & Nielsen 2009; Gravel 2012), but no competitors used them, perhaps due to insufficient outreach to the relevant subset of researchers. Methods for demographic history inference based on the Markovian coalescent (Li & Durbin 2011; Schiffels & Durbin 2014) that don’t depend on a user-specified parametric model were also underrepresented relative to their popularity in the literature.

The second GHIST competition launched at the Evolution conference in June 2025 in Athens, Georgia and runs through November 2025, with expanded challenge types and refinements based on lessons from the inaugural tournament. To discourage metagaming while preserving the benefits of iterative submission, for each challenge multiple submissions are allowed on a testing data set, but only a single submission is allowed on the final data set. To increase the realism and difficulty of demographic history inference, two of the challenges include background selection, leveraging the stdpopsim framework for simulation (Gower et al. 2025). To expand the range of tasks, four challenges involve inferring single or multiple hard selective sweeps (Stephan 2019), under simple and complex demographic scenarios and with and without background selection. The Synapse site for this second competition is at https://synapse.org/Synapse:syn65877330.

Ultimately, the success of GHIST depends on community participation. The more methods developers, users, educators, and students engage with the competitions, the more the community will learn. The space of potential challenges is vast, including inferences such as distributions of fitness effects (Eyre-Walker & Keightley 2007), spatial models (Bradburd & Ralph 2019), and polygenic selection (Barghi et al. 2020), and including complications such as lowpass data (Crawford & Lazzaro 2012), polyploidy (Dufresne et al. 2014), and biased gene conversion (Pouyet et al. 2018). Within Synapse, submissions are scored using custom code executed on a cloud instance, so more complex evaluation metrics are possible. This initial competition has demonstrated the feasibility and utility of competitions for this community. Future competitions will enable deep insight into best practices for population genetics inference.

## Acknowledgments

This work was supported by the National Institute of General Medical Sciences of the National Institutes of Health (R35GM149235 to R.N.G.). A.D. received support from NIGMS predoctoral training grant 5T32 GM067553. We thank Pablo Meyer Rojas for introducing us to the Synapse platform and the Society for Molecular Biology and Evolution for hosting the kickoff workshop and promoting the competition on their social media channels. We thank all competitors for their participation.

This paper was written with the assistance of generative artificial intelligence (AI). MacWhisper was used to transcribe the audio from author Gutenkunst’s talk about GHIST at the 2024 Probabilistic Modeling in Genomics conference. Anthropic’s Claude 4 Sonnet model was then given that transcript and asked to generate a detailed outline for the paper, including leveraging its existing knowledge base, within a Project containing previous papers by author Gutenkunst. Author Gutenkunst also used Claude’s research mode to generate a report on published papers that independently compared population genetics inference approaches, which yielded a few studies he was previously unaware of. Author Gutenkunst edited or generated all text in the final manuscript and verified all references.

